# Integration of imaging modalities with lipidomic characterization to investigate MSCs potency metrics

**DOI:** 10.1101/2022.05.25.493259

**Authors:** Priyanka Priyadarshani, Alexandria Van Grouw, Adrian Ross Liversage, Arina Nikitina, Kayvan Forouhesh Tehrani, Melissa Kemp, Facundo Fernandez, Luke J. Mortensen

## Abstract

Mesenchymal stem cells (MSCs) are widely used as therapeutics targets for numerous autoimmune diseases. However, MSC therapies have had limited success so far in clinical trials, mainly being heterogenous population it is difficult to determine MSCs efficiencies. It is critical to understand internal signaling of individual MSCs population that directly affect the cell phenotype. Lipid signaling is closely associated with cell shape so, a holistic approach to understand how changes in lipid metabolites trickles all the way to single cell phenotype could reveal deeper understanding of MSCs functional regulation. So, we aim to evaluate lipid metabolic profiles of single cell MSCs with known variability in immune regulation and explore the phenotypic changes that occur because of differences in lipid signaling. We use longitudinal label free phase imaging strategies to obtain cell phenotypic features which are directly correlated with single cell lipid metabolome obtained using advanced MALDI-MSI technique. Correlation maps indicate associations between lipid signaling and phenotypic changes in MSCs.

Moreover, a novel machine learning clustering approach detects the heterogeneity in the MSCs subpopulation then methodically see how each heterogenous population is being impacted by the changes in lipid profiles which could be linked to the functional behaviors of the cell.

## Introduction

Mesenchymal stromal cells (MSCs) offer great therapeutic potential for the treatment of autoimmune and inflammatory diseases^1,2^. MSCs are gaining scientific interest for their clinical use, however, phenotypic and functional heterogeneity in MSCs limits the therapeutic efficacy and complicates the use of MSCs in regenerative applications^3,4^. Although MSCs are treated with inflammatory cytokines such as interferon-gamma (IFN-γ) to enhance their functional properties, not all cells become equally active^5^, and thus there is a difference in the functional behavior within the subpopulation of MSCs. So, it is critical to characterize the functionally active subpopulation of MSCs. A study for single-cell analysis to closely monitor the cell signaling and change in morphological phenotypes can yield further insights into the mechanisms and implications of single-cell variation amongst these cells and can help enhance their clinical efficacy. There is growing evidence that lipids play a vital role in MSC functions and are closely associated with the MSCs morphological phenotype^6,7^. Although lipid metabolism plays a pivotal role in MSCs physiopathology, the number of studies at the moment about the lipidome of MSCs is limited and no strong link has been established to understand the direct relationships between single-cell MSC lipidome and its phenotype. We developed a study pipeline to simultaneously obtain and link the phenotypic features and lipidomic profiles of single-cell MSCs from the same monolayer culture using two label-free imaging techniques, a quantitative differential phase contrast (DPC) microscope to obtain phenotypic features and matrix-assisted laser desorption/ionization (MALDI) mass spectrometry imaging (MSI) to obtain a lipidomic profile. Using these methods, we show the association between changes in lipidomic profiles and the simultaneous change in phenotypic features of IFN-γ stimulated MSCs. Additionally, we identify morphological features that were associated with functional lipid signaling, mostly involved in the biological process of MSC functions. These findings highlight the close association between lipid and morphology and can be used as a predictive marker to filter the functionally active subpopulation of MSCs and improve the MSCs’ quality for regenerative applications.

## Results

### 1. Label-free imaging shows heterogeneity and a shift in MSCs morphology with IFN-γ stimulation

Our main goal was to perform simultaneous morphological and lipidomic characterization of MSCs on the same set of cells with the ability to track features at the single-cell level. In order to achieve this, we first adopted a label-free non-invasive technique to obtain morphological features through whole-well images using a DPC microscope system. MSCs were first seeded on Indium Tin Oxide (ITO) coated slide. Half the wells in each ITO slide were stimulated with IFN-γ while the other half was left untreated as a control (Fig 1A). All the cells were then used to obtain whole-well images using a DPC microscope system (Fig 2B, 2C). The advantages of this cell imaging system were four-fold: First, the non-invasive method ensured that we avoided irreversible changes or destruction of the sample cells. Second, the collection of whole well images allowed us to track results for individual cells through their spatial arrangement and overlay them with images obtained through MALDI-MSI. Thirdly, using a label-free method ensured we capture our results with no alterations. And finally, the ITO-coated slides used to seed the cells in this study are compatible with both DPC and MALDI-MSI imaging techniques.

**Fig 1.**
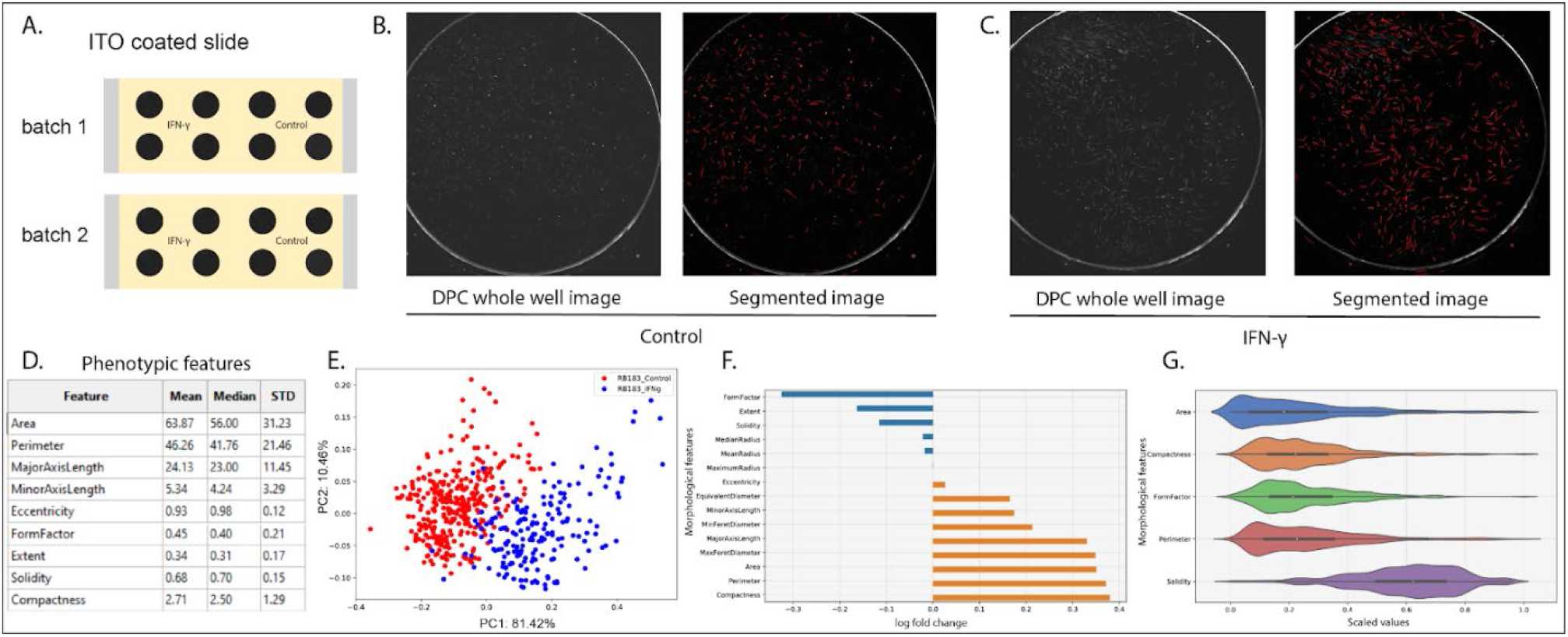
1. Label-free imaging shows heterogeneity and a shift in MSCs morphology with IFN-γ stimulation. Label-free DPC whole well imaging and cell segmentation. MSCs seeded on ITO-coated slides. Half of the well treated with IFN-γ and half of the cell untreated control (1A). Whole well DPC images of MSCs unstimulated (1B, left) and stimulated with IFNγ (1C, left). Cell segmentation of using cell profiler. The segmented unstimulated cells (1B, right) and stimulated cells (1C, right) are outlined in red. List of morphological features obtained from the cell profiler for each individual cell (1D). PCA plot showing separation between morphological features of stimulated and unstimulated MSCs (1E). Chart showing changes in morphological features after IFN-γ stimulation (1F).Volcano plot showing the distribution of shape within the subpopulation of MSCs (1G).

**Fig 2.**
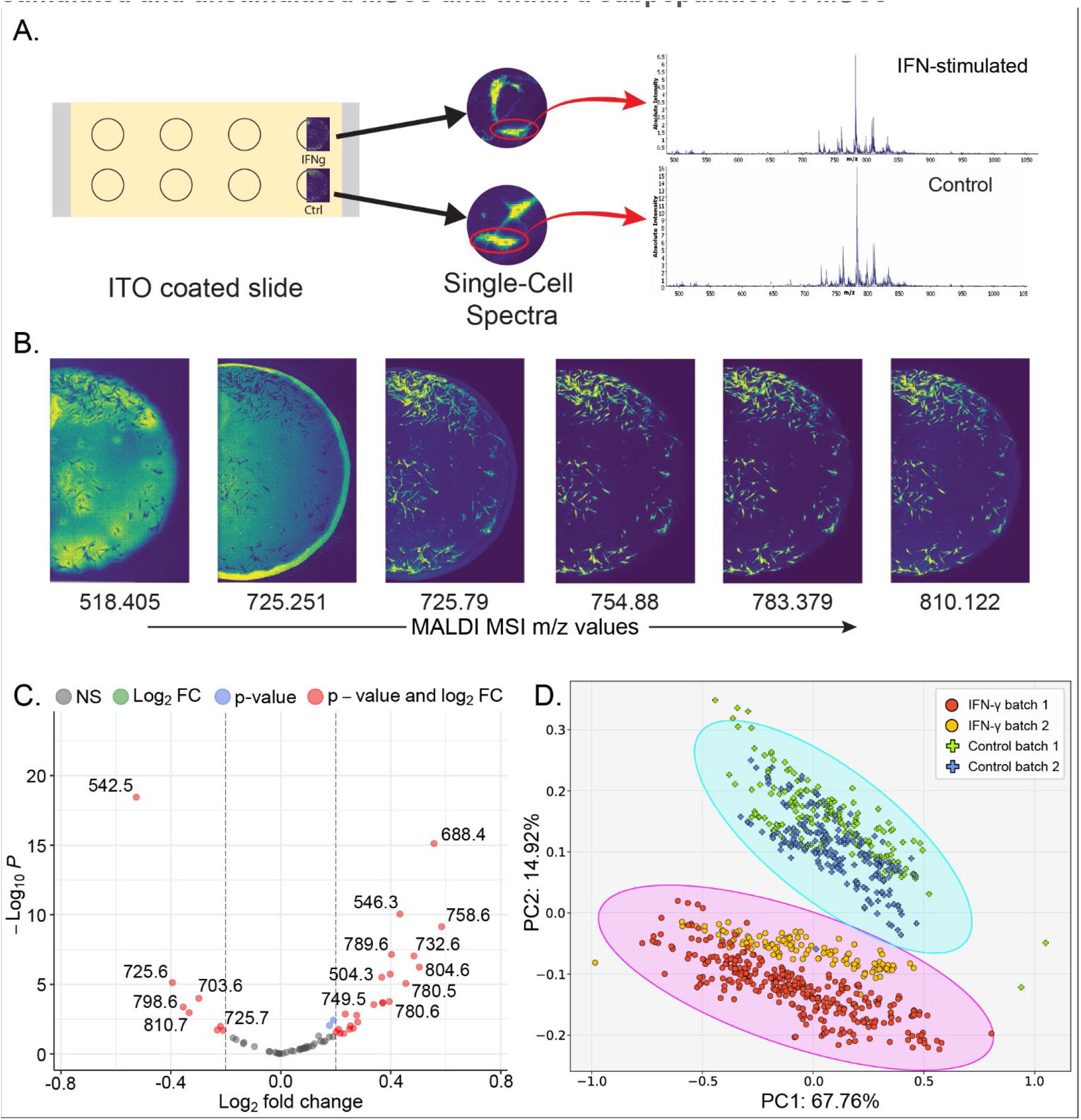
2. MALDI-MSI imaging of lipid peaks shows differences in lipid intensity between stimulated and unstimulated MSCs and within a subpopulation of MSCs. Single-cell lipid detection. MALDI imaging technique showing the detection of lipid peak spectra in single-cell stimulated and unstimulated MSCs (2A). Representative MALDI images of lipid peak for each individual MSCs (2B). Volcano plot showing lipid peaks higher in stimulated and unstimulated MSCs (2C). PCA plot showing separation between stimulated and unstimulated MSCs based on MALDI-MSI data (2D).

The obtained whole well DPC images were segmented using a cell-profiler (ref) pipeline (see Methods) designed to identify cell boundaries and extract cell exterior features (Fig 1B, 1C). Overall, 16 morphological features including cell phase intensities of both stimulated and unstimulated single-cell MSCs were obtained (Fig 1D) (supplementary table 1). As expected, Principal Component Analysis (PCA) showed separation between INF-γ stimulated and unstimulated MSCs. A total of 92% variance was explained by the first two principal components (Fig 1E). The bar chart in (Fig 1F) summarizes how the MSC morphological features changed with INF-γ stimulation. Some of the morphological features are increasing while others are decreasing. Overall, using our label-free DPC imaging technique and segmentation, the trends in morphological changes of MSCs upon IFN-γ stimulation were found to be consistent with previous observations^8^. We also note that the observed response to INF-γ stimulation is quite heterogeneous in shape. Within MSCs subpopulations after INF-γ treatment showed that not all cells are equally changing after stimulation (Fig 1G) highlighting the underlying caveat that not all cells are equally responding after stimulation.

### 2. MALDI-MSI imaging of lipid peaks shows differences in lipid intensity between stimulated and unstimulated MSCs and within a subpopulation of MSCs

The use of the label-free non-invasive method above allowed us to re-use the same slides for lipid characterization of MSCs at the single-cell level. The cell culture media was removed from the slides and air-dried so that the spatial location of the cells was preserved while fixing the cells without using any reagent or chemicals that might disrupt their molecular state and lipid content. The MALDI-MSI technique was used for the acquisition of images since it allows label-free imaging and does not require pre-treatment of the samples. MALDI-MSI data for individual MSCs were obtained in positive mode, using a smart beam laser setting of ~5 μm (average size of MSCs range from 10 to 20 um) and a laser raster size of 10 μm in both x and y dimensions to ensure sufficient spatial resolution. We were able to obtain both m/z peak spectra (Fig 2A) and a high spatial resolution image of each m/z peak (Fig 2B) (Supplementary Fig) for the single-cell MSCs. For each individual cell, a total of 66 images of lipid peaks were obtained. Cells within these images were colored individually based on the intensity of lipids detected. We noticed large differences in color intensities: both between the stimulated and unstimulated groups as well as within the same treatment group (Supplementary Fig) indicating a variegated response of cells to IFN-γ stimulation.

While differences in the IFN-γ treated group versus the control are expected^6^, it is the differences within each subpopulation that needs to be properly quantified in order to differentiate potent cells from unstimulated ones. To this end, we sought to measure the lipid proportions in each cell based on their coloring intensity using a cell profiler^9^based pipeline. The resulting cell-wise intensities were normalized before performing a comparison between IFN-γ stimulated versus control cells. The comparison revealed 25 lipid peaks that were higher in IFN-γ stimulated cells and 8 that were higher in the control cells. The remaining 34 peaks were unchanged between the two groups (Fig 2C) (Supplementary Table). In order to annotate the differentially expressed lipid peaks, we used popular online tools like Metascape (ref), and LipidMaps (ref) (Table 1). In addition, some peaks were also identified using Tandem Mass Spectrometry (MS/MS) (ref). We successfully mapped lipids to lipid group while peaks matched a group of lipids.

We next performed PCA on the matrix of differentially expressed lipid intensities in order to identify collective contributors to the separation of IFN-γ treated versus untreated cells. Each of the two treatment groups across two separate batches forms a tight group within the Hoetelling’s confidence ellipse indicating that there is no batch-to-batch variance in MALDI-MSI data after normalization. Overall, ~83% variance across both batches and treatments is explained by the first two principal components (Fig 2D). While variance within the subpopulation of MSCs is mostly explained by PC1, the separation between stimulated and unstimulated groups occurs along with the second principal component (PC2). This suggests that the main lipid profile contributors for the separation of stimulated cells from unstimulated cells are captured in PC2 (Fig 2D).

### 3. Coregistration of multimodal imaging modalities shows a correlation between MSCs morphological features and MALDI-MSI data in stimulated cells

In order to further understand what the contributors captured in PC2 are and how they contribute towards stimulation, we aimed to determine how the change in lipid signaling is related to the morphological changes. To establish the correlation between DPC and MALDI-MS images that we collected at the single-cell level we used co-registration pipieline. This pipeline took advantage of the intact spatial positioning of the cells across images generated from the two methods to computationally superimpose them by aligning and cropping to the same size (Fig 3A). Once this alignment was complete, it allowed us to use a unified cell-profiler pipeline that could identify, for every cell, the morphological features using the DPC image and then the lipid intensities using the series of MALDI-MSI images.

**Fig 3.**
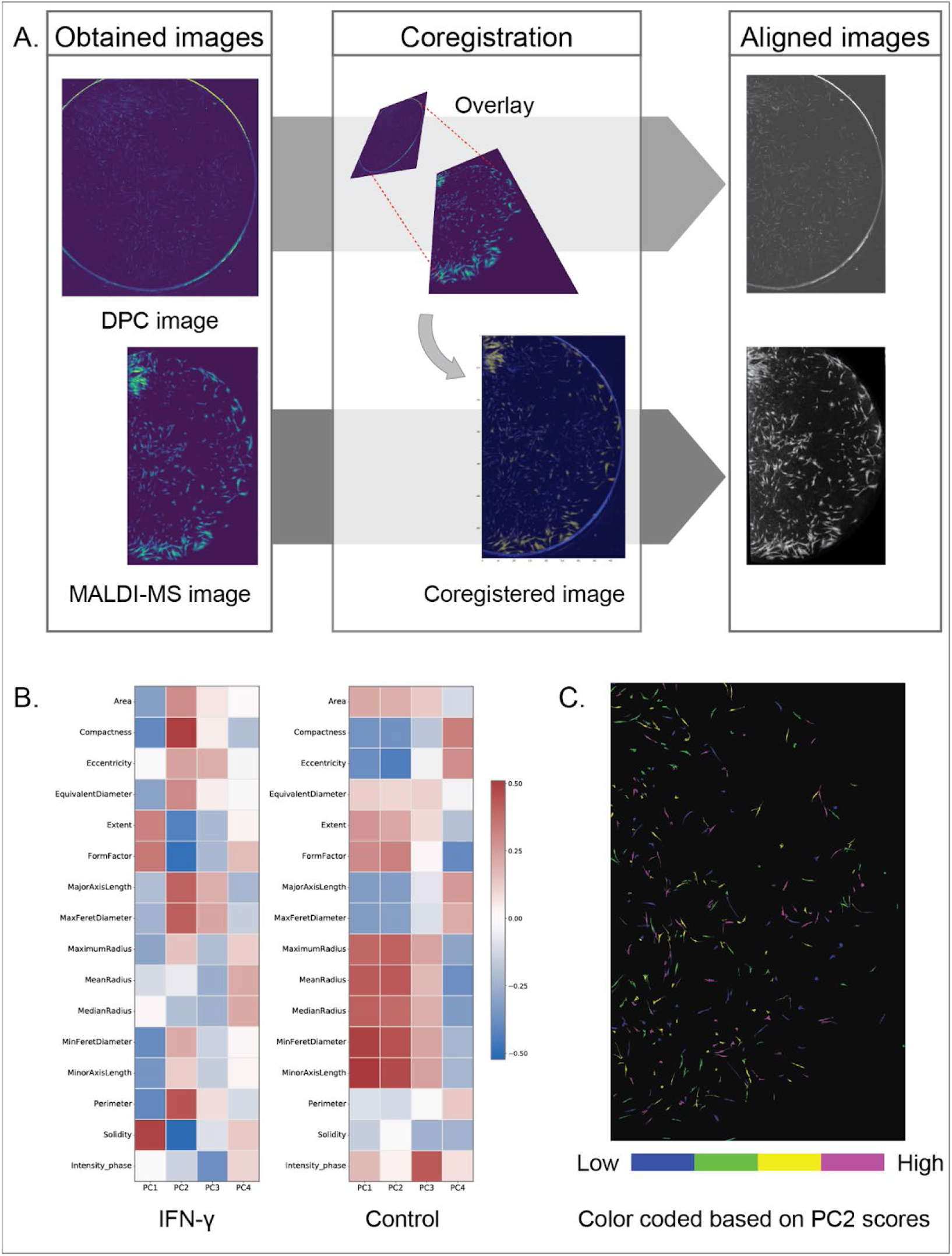
3. Coregistration of multimodal imaging modalities shows a correlation between MSCs morphological features and MALDI-MSI data in stimulated cells. Multimodal image coregistration. Coregistration of DPC and MALDI image (3A). Correlation map showing association of MSCs morphology with PC values based on lipid peaks in stimulated and unstimulated MSCs (3B). MSCs color-coded based on PC2 score obtained from MALDI-MSI data.

After this identification, in order to understand cell-to-cell relationships of morphological and lipid changes, Pearson’s correlation coefficient was calculated between the phenotypic features of MSC cells and their respective PC values obtained from MALDI-MSI data that captured the collective contributors of lipid intensities. In IFN-γ stimulated cells, MSCs morphological features like compactness, formfactor, perimeter, and solidity showed a high correlation with PC2 values. Interestingly we don’t see such a correlation in unstimulated cells (Fig 3B). This indicates that certain morphological features of MSCs are more responsive to lipid signaling after IFN-γ stimulation. To further confirm the above findings, we color-coded the MSC images with the range of PC2 values. The color-coding shows that not all cells have an equal intensity which confirms after treating the cells with IFN-γ, that not all cells are equally activated. These activated cells show certain morphological characteristics changes.

## Discussion

MSC-based cell therapy has gained great therapeutic interest and continued to be used at an alarming rate in pre-clinical and clinical trials despite showing inconsistent clinical effectiveness. The underpinning reason for this inconsistent clinical outcome is the heterogeneous nature of MSCs due to which their exact composition and regulatory mechanism are unclear. It is critical to understand the MSCs heterogeneity at the single-cell level and connect the internal signaling of each cell to its resulting external features which will contribute to establish the concise standards for MSCs cell-based therapy.

With this study for the first time, we were able to obtain a high spatial resolution image of the lipid peak of single-cell MSCs in monolayer culture and demonstrated the differences in lipid intensity within the stimulated and unstimulated groups and within the subpopulation of MSCs. We identified highly expressed lipids after IFN-γ stimulation at the single-cell level which could be related to the MSCs functional mechanism. Moreover, by integrating the morphological features and lipidomic profile of single cell MSCs make it possible to identify the phenotype associated with the functional lipid in MSCs. This could open the possibility to target and filter the cell based on phenotype for enhanced and consistent clinical therapeutics.

## Methods

### Cell culture

Human Mesenchymal Stem cells (hMSCs) (RoosterBio) were seeded on ITO coated slide (Millipore Sigma) under standard culture conditions. Culturewell gasket of well size 6mm diameter and 1mm depth (Grace Bio-labs) was used to on top of ITO slide to hold cells and media. Cells were seeded at a density of 1200 cells per well. The cells were grown in MEM alpha medium (Thermo Fisher Scientific) supplemented with 10% fetal bovine serum (FBS), 1% penicillin/streptomycin, and 0.5% of L-glutamine. The cultured cells were incubated with 5% CO2 at 370C for 24 hours. After 24 hours of incubation, either the regular cell culture media was replaced by conditioned media having 50 ng/ml IFN-γ or just replaced with fresh regular culture media. The MSCs under both conditions were incubated for an additional 24 hours. After 24 hours of incubation the cell were taken for imaging.

### Cell imaging

The images were sourced using an illumination-based differential phase contrast (DPC) microscope system as described in^10–12^. The illumination source is a 32×32 (4mm pitch) RGB LED matrix panel (product ID 607, Adafruit) controlled by an Arduino Uno (Arduino.cc) with LEDs set to hex color #1AFF00, which provides light of a broad spectrum around the suggested wavelength of 514nm. A 10x objective lens (Plan Fluor, Nikon) placed in the chassis of an inverted microscope collects the light relaying the image onto the sCMOS camera (Zyla 4.2, Andor) which is conjugated to the sample plane.

### Whole well DPC image construction, segmentation, and data analysis

Individual DPC images with a field of view of 1.3mm2 are captured sequentially, as an X-Y stage is automatically moved in steps of 1mm, capturing 49 sub-images (7×7), accounting for the whole circular well with 8mm diameter. Once the DPC reconstruction of each sub-image has been completed, the superfluous 300μm2 of each sub-image is removed. Using the built-in “Make Montage” function in Image-J, these sub-images are stitched together appropriately, representing the whole well image.

An image analysis pipeline was established in CellProfiler 4.0.7 was applied to whole well cell DPC images to extract morphological features consisting of 15 morphological features and a phase intensity feature.

Overall shift in MSCs morphology upon IFN-γ stimulation was determined by performing PCA on the 16 morphological features of the MSCs obtained from cell profiler. Log fold change was calculated for all the 16 morphological features to determine the magnitude of their changes. To illustrate heterogeneity within each of the morphological features, the scaled values were plotted as violin plots. All the analysis were done using custom python script.

### MALDI Imaging and MALDI data collection

MALDI-MSI^13^ was performed in positive mode on a RapifleX Tissuetyper time-of-flight mass spectrometer (Bruker Daltonics) equipped with a Smartbeam3D 10 kHz Nd:YAG (355 nm) laser. To control imaging settings, FlexImaging 4.0 software was used. To control pixel size, the smart beam laser setting (~5 μm in both x and y dimensions) with a laser raster size of 10 μm in both x and y dimensions was selected. Mass calibration was performed prior to data collection with red phosphorus as the calibrant. Additional experiments were performed using an ultra-high resolution Bruker solariX MALDI FTICR^14^ (Bruker Daltonics) with internal calibration using the exact masses of two known peaks. MALDI laser settings consisted of 25% power with medium focus and 25 μm raster width. Data were collected at a 1M transient size with range 300-1200 *m/z*. Preprocessing of the mass spectral data was performed in SCiLS Lab software. Baseline removal was performed during data import using iterative convolution and Root-Mean-Square (RMS) normalization was done in SCiLS. MSI images in SCiLS were exported in imzML and ibd formats for further processing. SCiLS lab was used to isolate lipid spectra of single cells using the polygon tool. The peaks present in cells only were used to create a peak list.

### Multimodal Image co-registration

The images obtained from DPC and MALDI imaging were aligned using co-registration pipeline developed by Nikitina et. al using custom python script^15^. Parameters within the pipeline were optimized to establish co-registration between the DPC and MALDI-MS images. Having the prior information of image sizes from both imaging system, with pipeline the images were scaled, shifted coordinately, and scaled until the best alignment is established between the images. After alignment, the images were cropped by alignment matching region. The location and region of each cell were matched in both images.

### Data analysis and Statistical Comparison

Once the location of each cell from both DPC and MALDI images were matched, CellProfiler pipeline (Supplementry) was used to extract the morphological features from the DPC images and the lipid intensity data at each m/z value on a cell from MALDI images. The lipid intensity at each m/z value was normalized by Standard deviation and scaled using min-max scaler. Student’s t-test was performed to determine the highly expressed lipids in IFN-γ stimulated cells. A cutoff of log fold change > 0.3 and P-values <0.05 was used to identify lipids that were significantly differentially expressed between the control and stimulated cells. PCA was performed using these significant lipid intensities across all 4 batches (2 control and 2 stimulated) to determine consistency across batches. The top 4 principal components (PCs) were extracted that explained nearly 91% of variability in the data. For each of these 4 PCs, correlation was calculated against all morphological features from control and stimulated cells separately. Since PC2 produced the most separation between control and stimulated cells in PCA, the PC2 scores were used to color code the cells using custom python scripts. The highly differentiated peaks were annotated using Metaspace and LIPID MAPS matches (Table1).

